# Engineering gene overlaps to sustain genetic constructs in vivo

**DOI:** 10.1101/659243

**Authors:** Antoine L. Decrulle, Antoine Frenoy, Thomas A. Meiller-Legrand, Aude Bernheim, Chantal Lotton, Arnaud Gutierrez, Ariel B. Lindner

**Author notes:** co-first authors (Antoine L. Decrulle, Antoine Frenoy).

## Abstract

Evolution is often an obstacle to the engineering of stable biological systems due to the selection of mutations inactivating costly gene circuits. Gene overlaps induce important constraints on sequences and their evolution. We show that these constraints can be harnessed to increase the stability of synthetic circuits by purging loss-of-function mutations. We combine computational and synthetic biology approaches to rationally design an overlapping reading frame expressing an essential gene within an existing gene to protect. Our algorithm succeeded in creating overlapping reading frames in 80% of *E. coli* genes. Experimentally, scoring mutations in both genes of such overlapping construct, we found that a significant fraction of mutations impacting the gene to protect have a deleterious effect on the essential gene. Such an overlap thus protects a costly gene from removal by natural selection by associating the benefit of this removal with a larger or even lethal cost. In our synthetic constructs, the overlap converts many of the possible mutants into evolutionary dead-ends, effectively changing the fitness landscape and reducing the evolutionary potential of the system.

## Introduction

Synthetic biology attempts to use engineering principles to manipulate and reprogram living organisms [6]. This could hold promise for many of the world’s challenges, for example with microbes engineered for bioremediation [3, 23] and drugs or fuel biosynthesis [26, 2]). Rational genetic engineering poses many design constraints, that are dealt with approaches stemming from electrical engineering. However, evolution brings a particular set of challenges [5, 13].

Synthetic systems are generally costly to their hosts, and mutants that alter or neutralize them will eventually be selected (figure 1a). Beyond inactivation by mutations, unforeseen evolution of circuits released in the wild also raises strong concerns. Much research effort is thus dedicated to the restriction of evolutionary potential [34, 41, 29, 30]. Similarly, containment and control of engineered organisms outside of the laboratory relies on sophisticated gene circuits that should be made as evolutionary-proof as possible [16, 8, 37].

**Figure 1:**
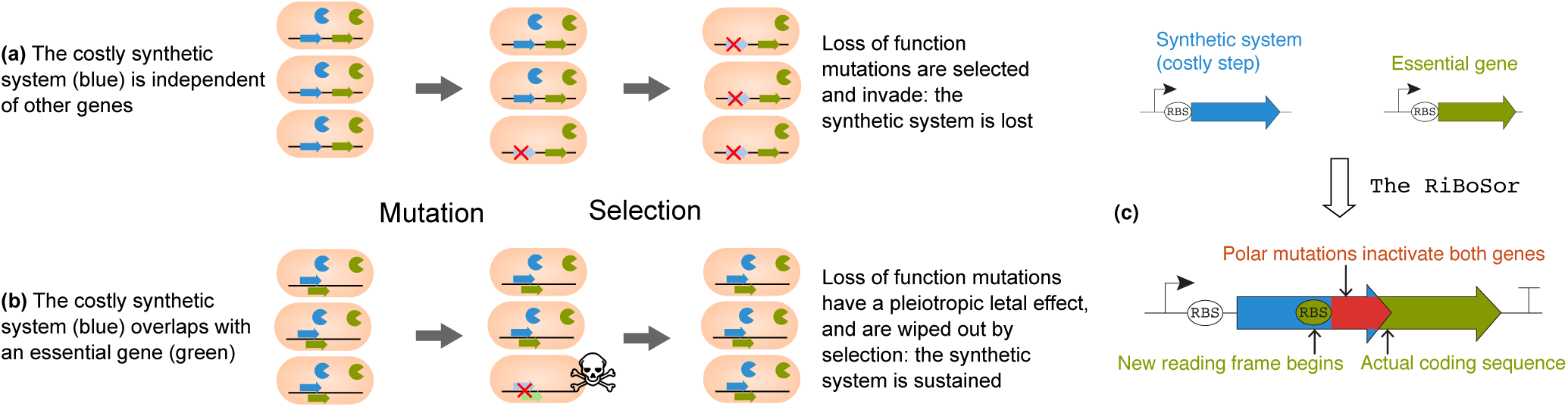
Principle and construction of a Riboverlap. **(a)** Standard case: loss-of-function mutations in the synthetic circuits are selected, because they alleviate the cost of the system. **(b)** The synthetic gene overlaps with an essential gene: loss-of-function mutations are discarded by natural selection because they induce a lethal cost. **(c)** Rational design of a Riboverlap: a new reading frame is created within the gene to protect (blue). An essential gene (green) is cloned downstream, within this reading frame. The overlapping part (red) is fused on the 5’ end of this essential protein.

We recently postulated that in nature, coding sequences can evolve overlapping reading frames as a way of reducing their evolvability [15]. Such gene overlaps impose strong constraints on sequences and their evolution [27, 20, 33, 21]. If a synthetic gene of interest overlaps with an essential gene, many of the loss-of-function mutations will also affect this essential gene and be evolutionary dead-ends (figure 1b). This led us and others [5] to suggest that gene overlaps could be engineered and used as a method for preventing gene loss in synthetically engineered organisms.

In this work, we test the use of overlapping reading frames to protect a gene from mutations. We first designed an algorithm to create a new reading frame within an arbitrary coding sequence, without modifying the encoded protein. Secondly, we experimentally assessed the evolutionary trajectory of the obtained synthetic constructs by quantifying loss-of-function mutations. We found that overlapping reading frames can be constructed in a large fraction of genes, spanning all bacteria and all functional categories; and that they bring a significant protection from mutations.

These results show that evolutionary constraints can be harnessed to enhance the robustness of systems costly to their host organism. We unveil a promising method to do so, and release our software implementation, the RiBoSor, under a GPL license permitting use and modification by the community.

## Results

### Algorithmic design of Riboverlaps

We developed an algorithm, the RiBoSor, to design an overlapping reading frame within an existing coding sequence (figures 1c, S1). This is achieved by creating a Ribosome Binding Site (RBS) followed by a start codon within the coding sequence, in a different reading frame. The sequence is then further modified to make the newly created reading frame suitable for expression of a protein. For example stop codons in the new reading frame are removed with substitutions that are synonymous in the existing reading frame.

An essential gene is cloned downstream the existing sequence, in the newly created reading frame (figure 1c). The essential protein is thus N-terminally fused with the end of the original sequence translated in another reading frame.

The efficiency of our design (called hereafter Riboverlap) stems from the protection it confers against polar mutations (*e.g.* large rearrangements, most indels, and IS transpositions). While the overlapped portion of our construct is not *stricto sensu* coding for the essential gene, polar mutations located in this portion will also affect this gene and be purged by natural selection. A large fraction of loss-of-function mutations are thus removed from the mutational pool due to the pleiotropy induced by the Riboverlap.

### Theoretical protective effect of Riboverlaps

The strength of the protection depends on the size of the Riboverlap, the proportion of mutations that are frameshifts, and the expected effect of a frameshift mutation compared to the effect of a base pair substitution. We study the effect of these parameters using Monte-Carlo simulations of mutations in a Riboverlap (figures 2a, S2). The parameter range was chosen based on the literature (Methods). We find that it is possible to obtain a protection of up to 80% of mutation removal for an entire Riboverlap in a wild-type strain, with realistic parameter values.

**Figure 2:**
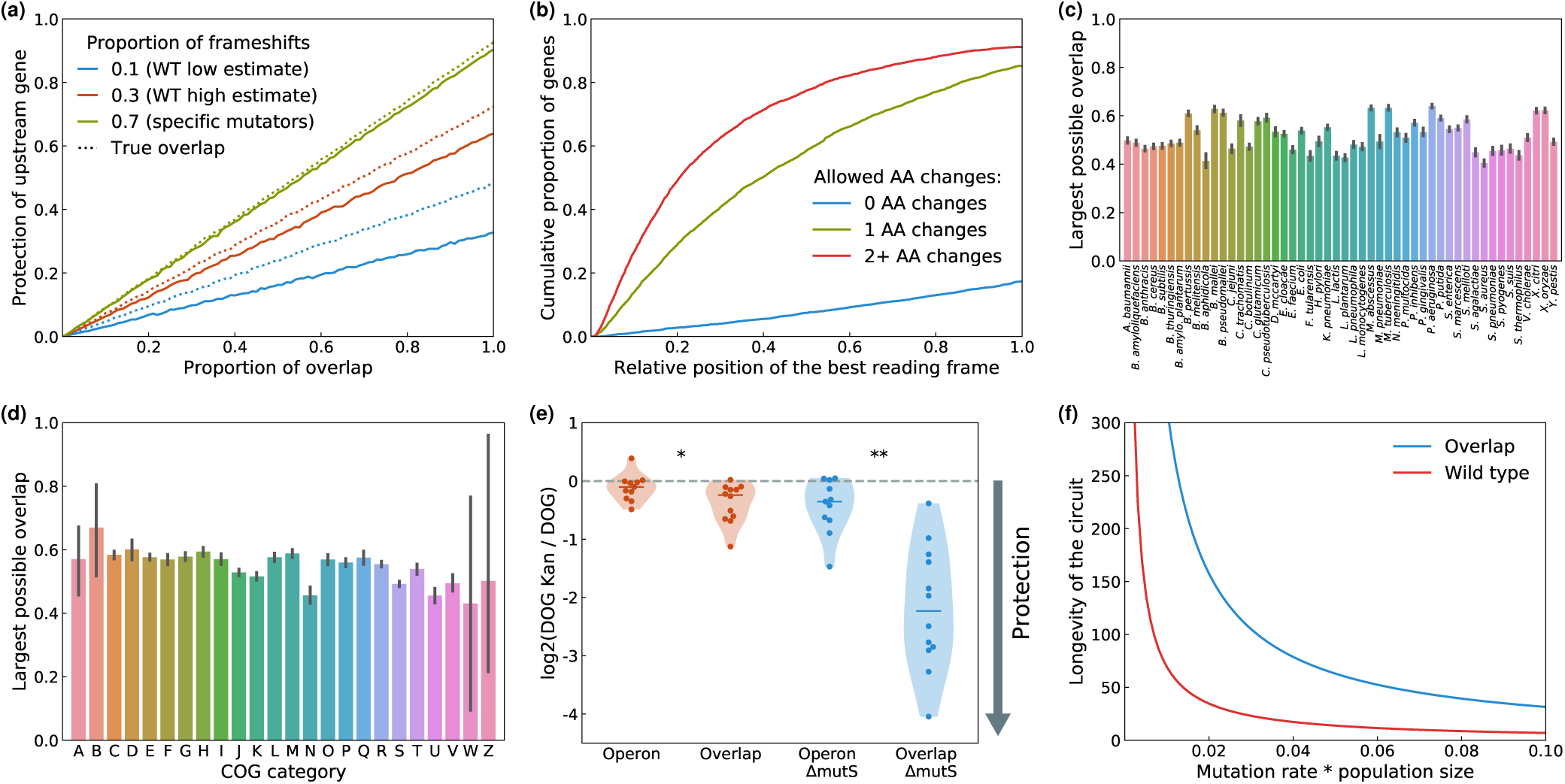
Our algorithm can create large overlapping reading frames in many genes spanning all organisms and functional categories. **(a)** Simulated protection conferred by a gene overlap, depending on the size of the overlap and the fraction of frameshift mutations. A true protein overlap (dashed lines) would also protect from non-polar mutations. **(b)** Potential Riboverlaps in *E. coli*: cumulative distribution of the first position at which an overlapping reading frames can be created, for all genes of *E. coli* MGZ1. Different colors represent different levels of stringency: 0 (blue), 1 (green), or 2 (red) amino acid changes allowed. **(c)** Potential Riboverlaps in other bacterial species: relative size of the largest overlapping frame that can be created. All CDS of each organism are averaged, and intermediate level of stringency (1 AA change) was chosen. **(d)** Potential Riboverlaps in all COG categories. Pooling the 156,542 coding sequences of the 50 bacterial species, average relative size of the largest overlapping frame can be created, for each COG category. **(e)** The overlap protects galK from loss-of-function mutations. We measure the fraction of galK loss-of-functions (permitting growth on DOG) that did not affect kanR. The lower this fraction, the higher the protection. Each point represents an independent population. The violin plots show the median and kernel density estimation for 12 independent populations. **(f)** Theoretical median lifetime of the circuit, with the measured protection for the Δ*mutS* strain, depending on the rate of loss-of-function mutations and on population size.

### Constructing Riboverlaps in bacterial genes

We computed possible Riboverlaps within all coding sequences of *Escherichia coli* using the RiBoSor. The best protection is achieved by the largest overlaps, we thus consider the earliest position at which an overlapping reading frame can be created within each gene. We computed the cumulative distribution of this position, for three different stringency levels of our algorithm, allowing for no, one or several amino acid changes (figure 2b).

Building a Riboverlap without any amino acid change is only possible in 20% of *E. coli* genes. However, allowing a single change permits to create a Riboverlap in more than 80% of the genes, and a large part of these Riboverlaps (about 65%) have a size higher than half of the gene to be protected. Because gram-negative bacteria have a strong tolerance to non-canonical RBS [36] and because it is now possible to synthesize many variant constructs and test their phenotype, allowing one amino acid change is practically feasible.

We also tested the RiBoSor on 49 other model bacterial species, with a comparable success to that of *E. coli* (figures 2c, S3). Pooling all 156,542 coding sequences of the 50 bacterial species, we extracted the 41,177 coding sequences for which we were able to assign a COG category [38]. We found that it is possible to create Riboverlaps with comparable success for all COG categories (figure 2d).

### Construction of a Riboverlap *in vivo*

We experimentally tested our design using galK (galactokinase) as a candidate gene to protect. galK is costly in presence of DOG (2-Deoxy-D-galactose), a galactose analog (figure S4). The RiBoSor found 9 candidate Riboverlaps in galK, listed in table S5.

We synthesized a candidate Riboverlap that has a perfect consensus RBS and does not require any amino acid change. As the downstream essential gene, we used a kanamycin resistance gene (kanR) encoding the aminoglycoside phosphotransferase aph(3’)IIIa. As a control, we constructed a strain in which galK and kanR share the same promoter but without overlapping reading frame. Both the overlapped construct and the control (operon) could grow on kanamycin and showed sensitivity to DOG, demonstrating that both genes are functional and expressed (figure S6).

### Synthetic overlaps protect from mutations

We then test our hypothesis that the overlap with the essential gene (kanR) protects the costly gene (galK) from loss-of-function mutations. To do so, we quantify the fraction of galK spontaneous loss-of-function mutants that are also kanR loss-of-function mutants, by plating on DOG alone and DOG with kanamycin. If the overlap is protective, we expect a lower number of colonies on DOG with kanamycin than on DOG alone, as some galK mutants will also loose kanamycin resistance. We deployed our test in both wild-type and mismatch repair defective (Δ*mutS*) strains. The latter hypermutator not only has a significant frameshift-prone mutation spectrum [32], but is also relevant as often selected in long-term cultures [35].

As showed on figure 2e, in a wild-type *E. coli*, the overlap confers a small protection on the edge of statistical significance threshold (one-sided Mann-Whitney *U* test, *p* = 0.037, *U* = 95.0) compared to the operon. In a Δ*mutS* strain, the protection is stronger and highly significant (*p* = 1.8×10^−4^, *U* = 124.0), consistent with our simulations figure 2a.

Finally, we estimated the gain in temporal stability of the synthetic system expected from the measured decrease in mutation rate (Methods). We found that the temporal stability can be increased significantly in a large parameter range (figure 2f).

## Discussion

We show that the evolutionary constraints induced by gene overlaps can be harnessed to design evolutionary-robust synthetic systems.

Present in all domains of life, from phages [4] and bacteria [14, 12] to vertebrates [24] including mammals [39], overlapping genes present a fascinating puzzle. They were discovered in the first organism ever sequenced, the phage ΦX174 [4, 40], but the selective pressures and mechanisms leading to their evolution remained elusive for a long time. Some researchers found this phenomenon so bewildering that they interpreted it as an evidence that ΦX174 was engineered by an extraterrestrial intelligence using a very advanced DNA synthesis technology [42].

Subsequent hypotheses for the evolution of gene overlaps mostly involve genome compression, which can provide several benefits: better encapsulation [10], faster and cheaper replication of the genome [20], and smaller mutational load [11, 19].

Alternatively or as an additional selective force, we suggested [15] that gene overlaps could evolve as an evolvability suppression mechanisms, as defined by Altenberg [1]. In this work, we build upon this idea and show they can be used to restrict the evolutionary potential of synthetic systems.

Most of the field of synthetic biology relies on engineering-inspired modularity as a design feature.

The same way it is a constraint for sequence evolution, the lack of modularity induced by overlapping reading frames is generally seen as a hindrance for genetic engineering. This view prompted the refactoring of bacteriophages T7 and ΦX174 without gene overlaps [9, 18]. We follow an alternative approach, showing that a constrained, unmodular design can be an advantage in some situations and can be rationally designed and engineered.

Our design shares some conceptual similarity with the attempts to make promoters pleiotropic to protect them from downregulatory mutations [34, 41]. Here, beyond the promoter, we protect the coding sequence from the most deleterious classes of mutations: indels and transpositions of insertion sequences.

However, our method does not protect from nonpolar substitutions, implying that it will never provide perfect protection, as seen on figure 2a. Yet the protection is still substantial for large overlaps because the most deleterious mutations are frameshifts. Reducing the effective mutational supply can make a substantial difference in terms of effective stability of a synthetic system in a bioreactor as seen on figure 2f.

Our results provide a strong proof of concept that gene overlaps can be rationally engineered to reduce the evolutionary potential of synthetic constructs, that the absence of modularity can be a useful design feature, and that it is possible to rationally design constructs that do not only have a specific phenotype, but also particular evolutionary properties.

## Methods

### Simulation of the protection

We modeled protein loss-of-function resulting from mutations as a Bernoulli process, assuming that each amino acid change has a probability *P*_*e*_ of turning a fully functional gene into a non-functional gene. *P*_*e*_ can alternatively be understood as the average deleteriousness of each amino acid change.

Estimations of this parameter are classically reported based on mutagenesis data in the distribution of fitness effects literature. Most of this literature historically focused on RNA viruses, for which the estimates are relatively high and vary from 0.19 [31] or 0.37 [28] to 0.76 [7]. The estimates available for bacteria suggest that the value of *P*_*e*_ would be closer to the lower end of this range [25, 17]. We thus run sets of simulations for two different values of *P*_*e*_: 0.1 (figure 2a) and 0.3 (figure S2).

The other important parameter is *fs*, the proportion mutations that are frameshifts. Estimates in *Escherichia coli* vary between 0.1 and 0.4 for wild type strains, and up to 0.7 or 0.9 for specific frameshift prone mutators such as the Δ*mutS* mismatch repair defective mutant [22]. We thus explore the effect of three different values of *fs*: 0.1, 0.3 and 0.7.

For all combinations of the aforementioned parameters, we perform 100, 000 Monte-Carlo simulations for each possible size of the Riboverlap, defined as the proportion of the costly gene that is included in the Riboverlap. The output measure of our simulations is the fraction of the mutations impacting the costly gene that would be purged by natural selection because of their impact on the essential gene.

### Screening bacterial genomes

This RiBoSor was run with 3 different levels of stringency: 0, no non-synonymous changes allowed; 1, a single amino acid change allowed; or 2+, a single amino acid change allowed for the creation of the RBS-start, and as many as needed for the removal of the stop codons in the new reading frame – the rational being that there are many possible substitutions to remove a stop codon, and thus high chances that one of them is neutral although non-synonymous in the existing reading frame.

50 bacterial species (including *E. coli)* with the highest number of sequenced genomes in RefSeq were chosen for the screen. Computations were parallelized on an HPC platform.

### Strains and growth medium

Both constructs (overlap and control operon) were chromosomally integrated in the *intC* locus of *E. coli* MGZ1 Δ*galK*. The strain MGZ1 was constructed from MG1655 (CGSC #6300) by transduction (P1vir amplified on DH5alphaZ1) of the Z1 cassette, consisting of constitutively expressed copies of *lacI* and *tetR* and a spectinomycin resistance gene. The Δ*galK* deletion was obtained by P1 transduction from the Keio collection. The galK-kanR overlap and control operon were constructed using golden gate assembly into plasmids pOverlap and pControl, resp., next to a chloramphenicol cassette and flanked by 50bp *intC* homologies. MGZ1 was first transformed with the pKD46 recombineering helper plasmid and then with PacI-linearized DNA fragments of pOverlap and pControl. Δ*mutS* strains were obtained by P1 transduction of the Δ*mutS* allele from the lab stock.

Stock solutions of the following ingredients are prepared and autoclaved separately: ddH2O, M9×5, CaCl2 at 1M, MgSO4 at 1M, glycerol at 60%v/v, vitamin B1 at 0.1%*w/v*, agar at 30g/L. M9 glycerol medium is prepared by mixing 100mL of M9×5, 1.7mL vitamin B1, 1mL MgSO4, 50µL CaCl2, and 3.75mL glycerol with either 393.5mL ddH2O (liquid medium) or 143.5mL ddH2O and 250mL agar (solid medium, the agar is melted and cooled down at 56°C and the other ingredients are heated to 56°C before mixing). 2-Deoxy-D-galactose stock solution was prepared at 20%w/v and stored at 4°C, IPTG stock solution was prepared at 0.5M and stored at −20°C, and kanamycin stock solution was prepared at 100mg/mL and stored at −20°C.

### Phenotype of the constructs

Both the overlap strain and the operon (control) strain were able to grow in M9 glycerol supplemented with 25µg/mL of kanamycin, while the WT strain (*Escherichia coli* MGZ1 Δ*mutS*) was not. This confirms that kanR is functional (figure S6). In M9 glycerol supplemented with 0.2% of DOG, the overlap strain and the operon strain showed no growth after 24 hours at 37°C, while the WT strain could grow (figure S6). This confirms that galK is functional and can be used as a costly gene. The concentration of DOG we used was lethal in presence of galK, but could be decreased to modulate the cost.

### Mutagenesis protocol

N independent culture tubes containing 5mL of LB are inoculated with 5µL of a 10^−6^ dilution of an overnight culture (approximately 20 founder cells per culture). After 24 hours of growth at 37°C with vigorous shaking at 45° inclination, each culture is plated on two different solid media at appropriate dilutions: on M9 glycerol supplemented with DOG (final concentration 0.2%w/v) and IPTG (final concentration 0.5mM) to count galK loss-of-function mutants, and on the same medium with 25µg/mL kanamycin to count the fraction of these mutants that did not loose kanamycin resistance. The agar plates are incubated 64 hours at 37°C before counting colonies, due to slow growth on M9 glycerol medium. *N* = 11 for the control, and *N* = 12 for the overlap. The rank-biserial correlation *r*, calculated as 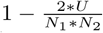, is *r* = −0.44 for the WT constructs and *r* = −0.88 for the Δ*mutS*constructs.

### Temporal stability of the circuit

We compute the median time to the first loss-of-function mutation in the costly system that does not inactivate the essential gene. This time is expressed in number of synchronous generations. Assuming the distribution of the number of mutational events in a given unit of time follows a Poisson law, the waiting time to the first event follows an exponential distribution.

### Data availability

All the experimental and theoretical data underlying this study, as well as the scripts used to analyze the data and produce the figures, are available on Zenodo (https://doi.org/10.5281/zenodo.3236452). The RiBoSor is available on GitHub (https://github.com/afrenoy/RiBoSor) and PyPi (https://pypi.org/project/RiBoSor).

## Acknowledgments

The authors thank Dusan Misevic, Edwin Wintermute, Tatiana Dimitriu and François Taddei for many discussions about gene overlaps, and Rebekka Wild for help with the graphical design of the figures. AD acknowledges financial support from the AXA Foundation PhD fellowship. ABL acknowledges financial support from the Fondation Bettencourt Schueller through the Center for Research and Interdisciplinarity.

## Supplementary material

The RiBoSor creates a translation initiation motif in the coding sequence of an input gene, and makes the new reading frame suitable for cloning and expression of a downstream gene.

The accuracy of our algorithm will depend on accurate prediction of translation initiation motifs. Existing thermodynamic models such as the RBS calculator are not tailored for the evaluation of RBS within coding sequences. Furthermore these tools are to slow to screen entire genomes, and their use is often restricted to slow web server without availability of the source code or a binary. We thus use a simple approximation: we consider that a translation initiation motif is a consensus Shine-Dalgarno sequence followed by 3 to 7 base pairs followed by a START codon. This simplistic approximation ignores important parameters such as secondary structure of messenger RNA. However, since our algorithm produces a library of possible constructs, it is possible to screen several candidates using a slower and more accurate model, or even to directly perform an experimental screen.

Considering all synonymous variants of a gene leads to a combinatorial explosion: for a 300 amino-acids sequence (typical *E. coli* gene), there are up to 3.2^300^ possible synonymous sequences (worst-case scenario with the average codon redundancy equally distributed, 3.2 being the average number of possible codons per amino-acid), which is well beyond what is computationally feasible. However, finding whether a subsequence can be rewritten to initiate translation is a local problem. We can thus brute-force smartly, by considering all the synonymous sub-sequences in a sliding window of an appropriate size. For each position in the input existing gene, the downstream 18 nucleotides (maximal size of the AGGAGG + spacer + START motif: 6+7+3, rounded to the next codon) are examined by a brute force algorithm. The algorithm probes every possible combination of synonymous changes and compares the resulting sequence to the target translation initiation motif. In the worst case, we examine 3.2^18^ possible sequences per position inside the input gene, reaching in total 300×3.2^18^ or about 3×10^9^ candidate sequences to evaluate, which is feasible in a reasonable time.

Different thresholds can be used to evaluate translation initiation motifs. Because gram-negative bacteria can use variants of the Shine-Dalgarno consensus sequence for translation initiation, we consider motifs that have up to one nucleotide substitution compared to this consensus.

Similarly, some amino-acid substitutions may be neutral, and it makes sense to allow for a small number of such non-synonymous changes. They may for example be needed to remove a stop codon in the new frame. Such removal can be achieved by many different substitutions, and the neutrality of every possible one can be tested experimentally.

The algorithm further analyzed candidate sequences that match the translation initiation motif, to introduce synonymous changes in the existing reading frame, to remove stop codons in the new reading frame, and, when possible, to optimize this new reading frame by removing other potential start codons and rare codons – which impede expression, and by removing mono-nucleotide repeats longer than three base pairs – which are mutagenic motifs.

**Figure S1:**
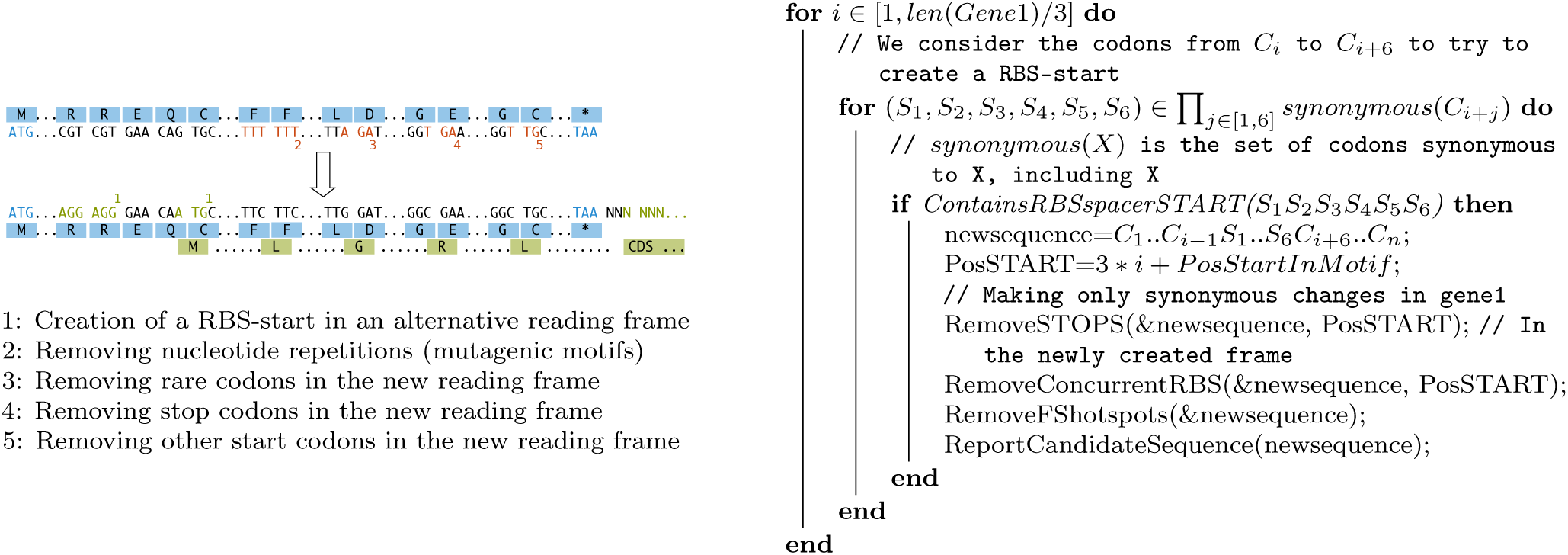
The RiBoSor, an algorithm to create alternative reading frame.

**Figure S2:**
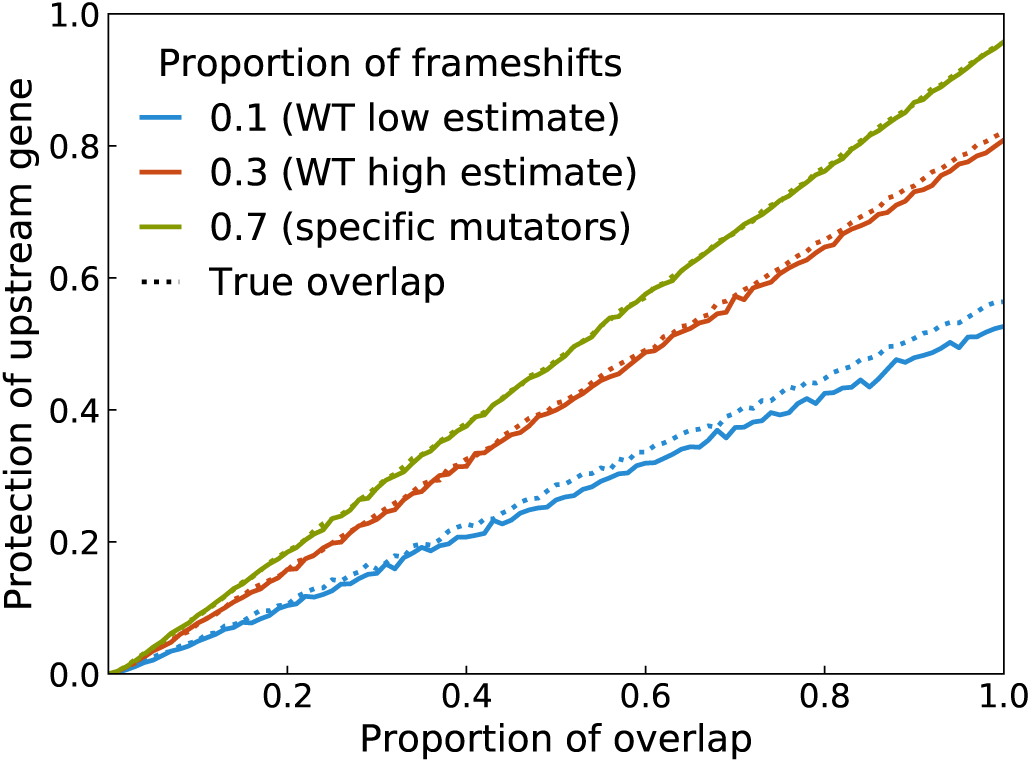
Simulated protection conferred by a gene overlap with. *P*_*e*_ = 0.3, depending on the size of the overlap and the fraction of mutations which are frameshifts. This is the same as figure 2a of main text, but instead of *P*_*e*_ = 0.1 (main text figure 2) that corresponds to a reasonable estimate for bacteria, we use *P*_*e*_ = 0.3, closer to what is observed for viruses.

**Figure S3:**
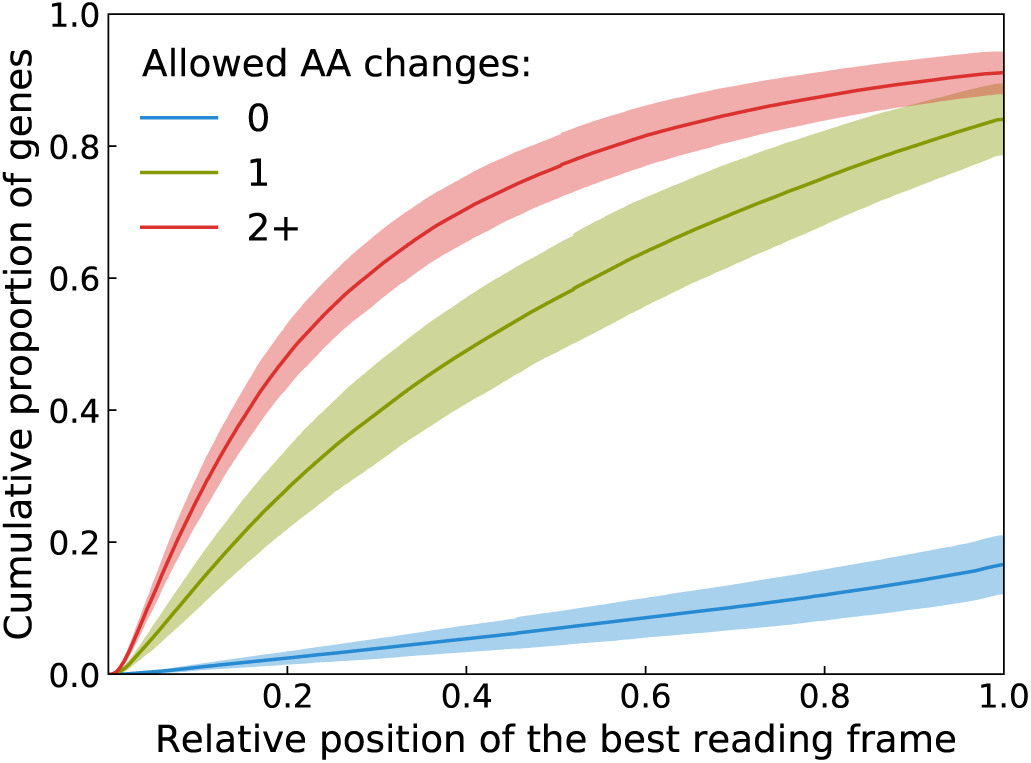
Potential riboverlaps in the genes of 50 representative bacterial species. Similarly to figure 2b, we run the RiBoSor on all genes of 50 representative bacteria species with the highest number of fully assembled genomes in NCBI database, as a proxy for their popularity in the microbial genomic research field. Full lines are the mean curves for the 50 bacterial species, and shaded area indicate the standard deviation.

**Figure S4:**
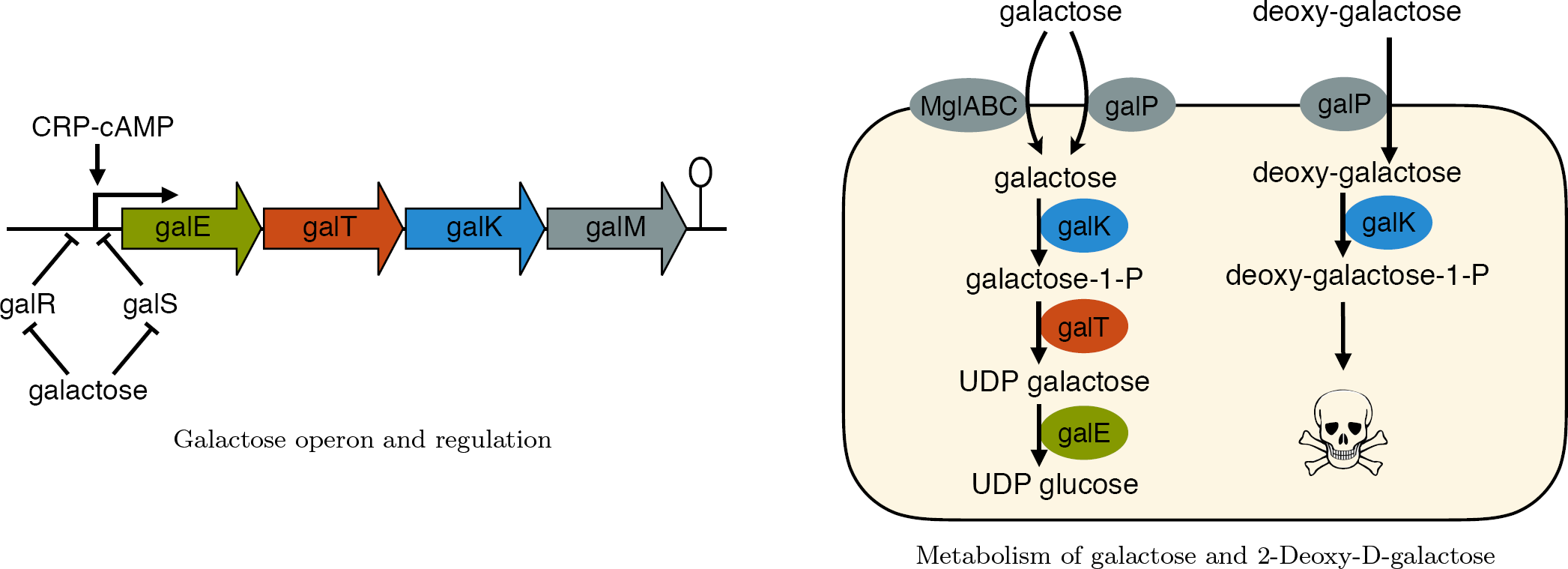
galK is a counter-selectable gene in presence of DOG. Galactokinase can be made costly and counter-selected by supplementing a non-glucose growth medium with 2-Deoxy-D-galactose (DOG). DOG is an analogue to galactose that can be imported by the same pathway and processed by galK, but can not be further processed by the rest of the galactose pathway and will accumulate into toxic intermediates. We used M9 glycerol as a carbon source to avoid the catabolite repression triggered by glucose. The expression of a functional galactokinase can be made visible without selection on Tetrazolium agar medium: the bacteria unable to ferment galactose appear as red colonies. galK can be positively selected (in galactose medium).

**Table S5:** Candidates found by the RiBoSor for galK. The algorithm indicates the number of synonymous and non synonymous changes made in the existing gene to create a new reading frame, and the remaining number of changes to be made manually by the experimenter. The bold line is the candidate chosen for experimental validation.

| Start pos. | Frame | Synonymous changes | Non synonymous | Remaining |
| --- | --- | --- | --- | --- |
| 475 | 1 | 96 | 1 | 1 |
| 544 | 1 | 91 | 1 | 1 |
| **675** | **2** | **74** | **0** | **0** |
| 742 | 1 | 75 | 1 | 1 |
| 995 | 1 | 61 | 1 | 0 |
| 998 | 1 | 62 | 1 | 0 |
| 1099 | 1 | 56 | 1 | 0 |
| 1129 | 2 | 31 | 1 | 0 |
| 1130 | 2 | 29 | 1 | 0 |

**Figure S6:**
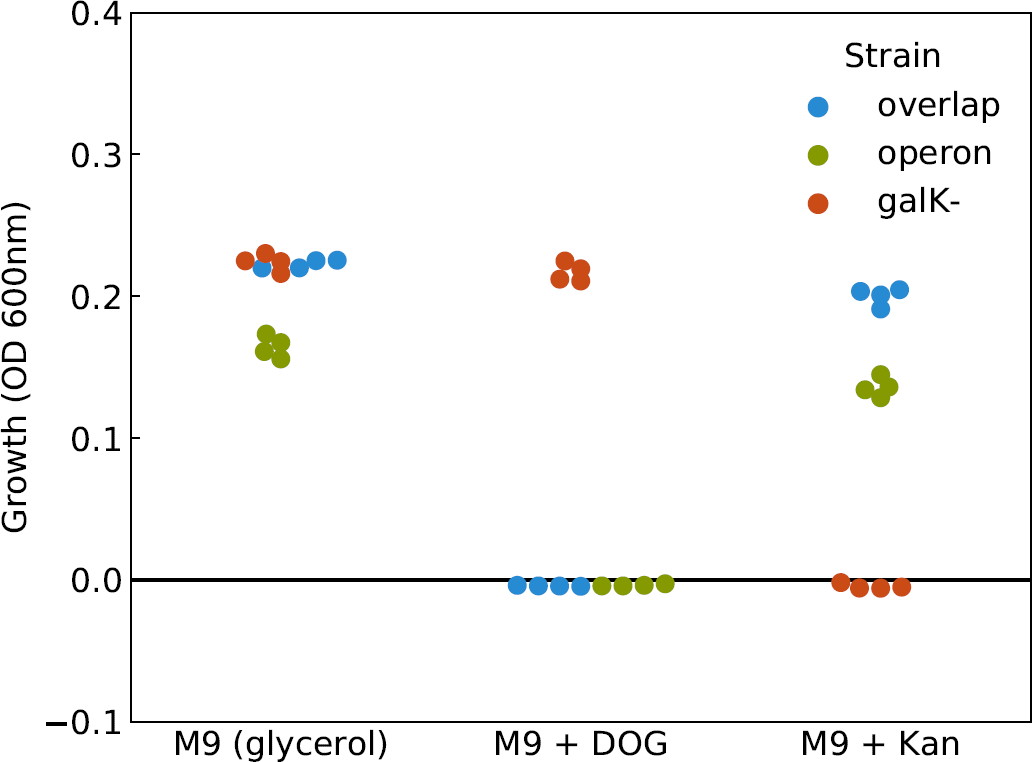
Phenotype of the galK-kanR synthetic overlap. In both the overlapping construct and the control (operon), galK and kanR are functional: the strains grow in presence of kanamycin and are inhibited by DOG.

